## Supplementary material for "Contrasting Effects of SARS-CoV-2 Vaccination vs. Infection on Antibody and TCR Repertoires"

### Figure S1: Timeline and sequencing yield

**(a)** Number of samples sequenced in each of several sequencing batches. **(b)-(c)** Repertoire sizes by immunosuppression status, cell type, and cohort.

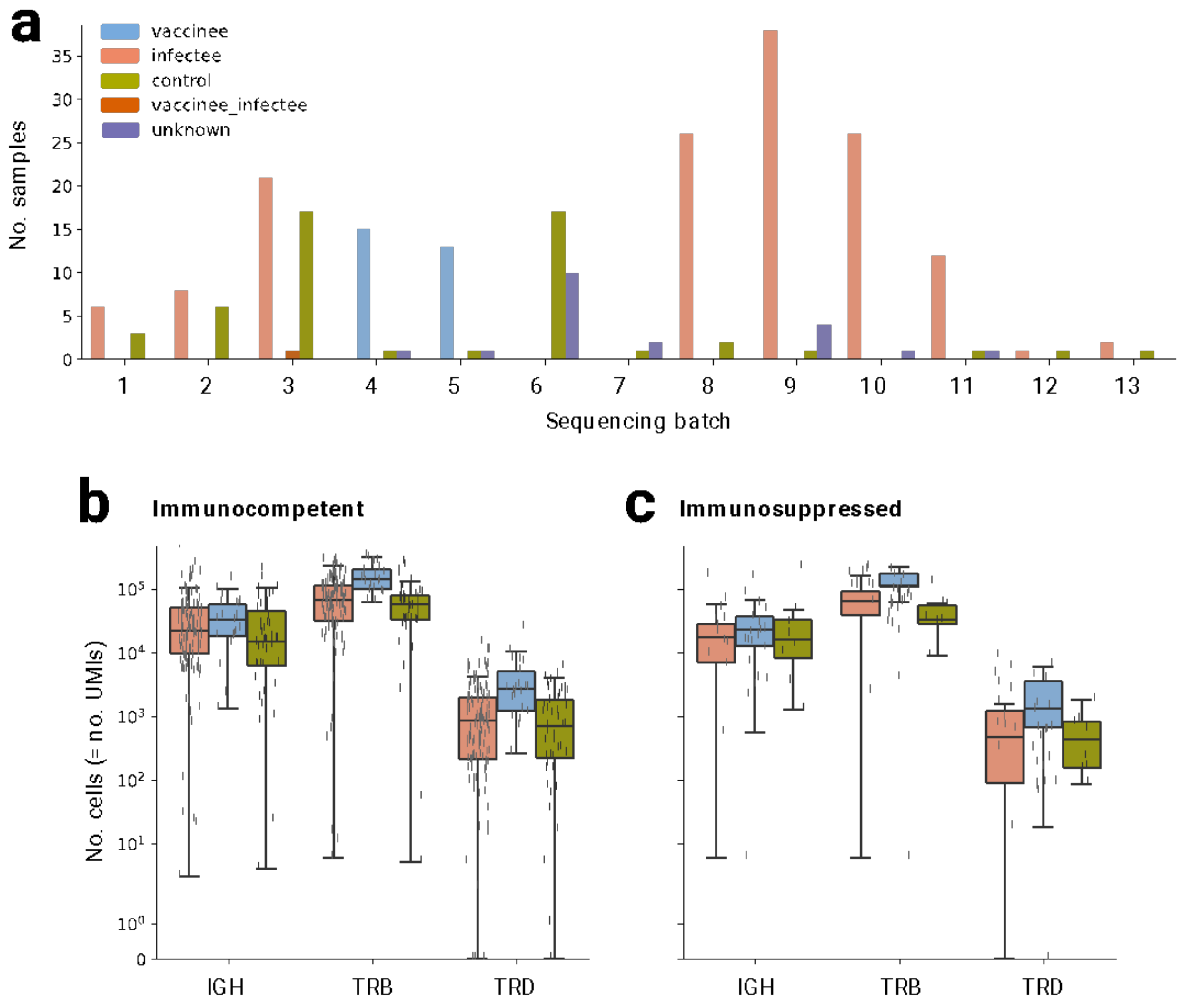

**Figure S2: CDR3 length comparison plots for functional IGH repertoire subsets**

Box-and-whisker plots of CDR3 length distributions (in nucleotides) in functional IGH genes in the control (olive) vs infectee (salmon) cohorts. (a) IgM-positive subset; (b) IgM-negative (predominantly IgG-positive) subset. See Fig. 2 legend.

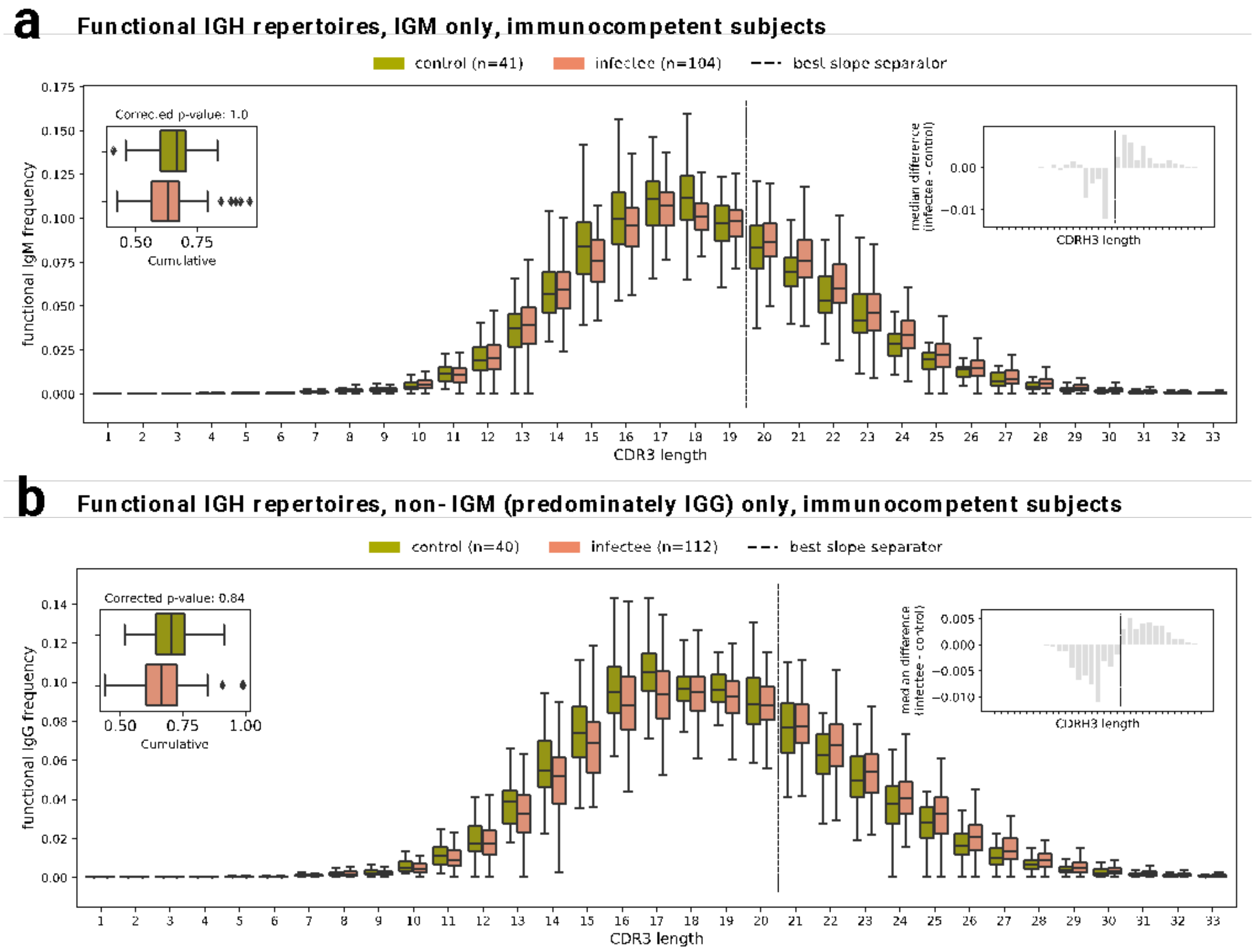

**Figure S3: CDR3 length comparison plots for functional TRB repertoires, including subsets**

Box-and-whisker plots of CDR3 length distributions (in nucleotides) in functional TCRβ genes. (a) Immunocompetent controls (olive) vs vaccinees (blue); (b) Immunocompetent infectees (salmon) vs vaccinees; (c) CD4-positive subset in immunocompetent controls vs infectees; (d) CD4-negative (predominantly CD8-positive) subset in immunocompetent controls vs infectees.

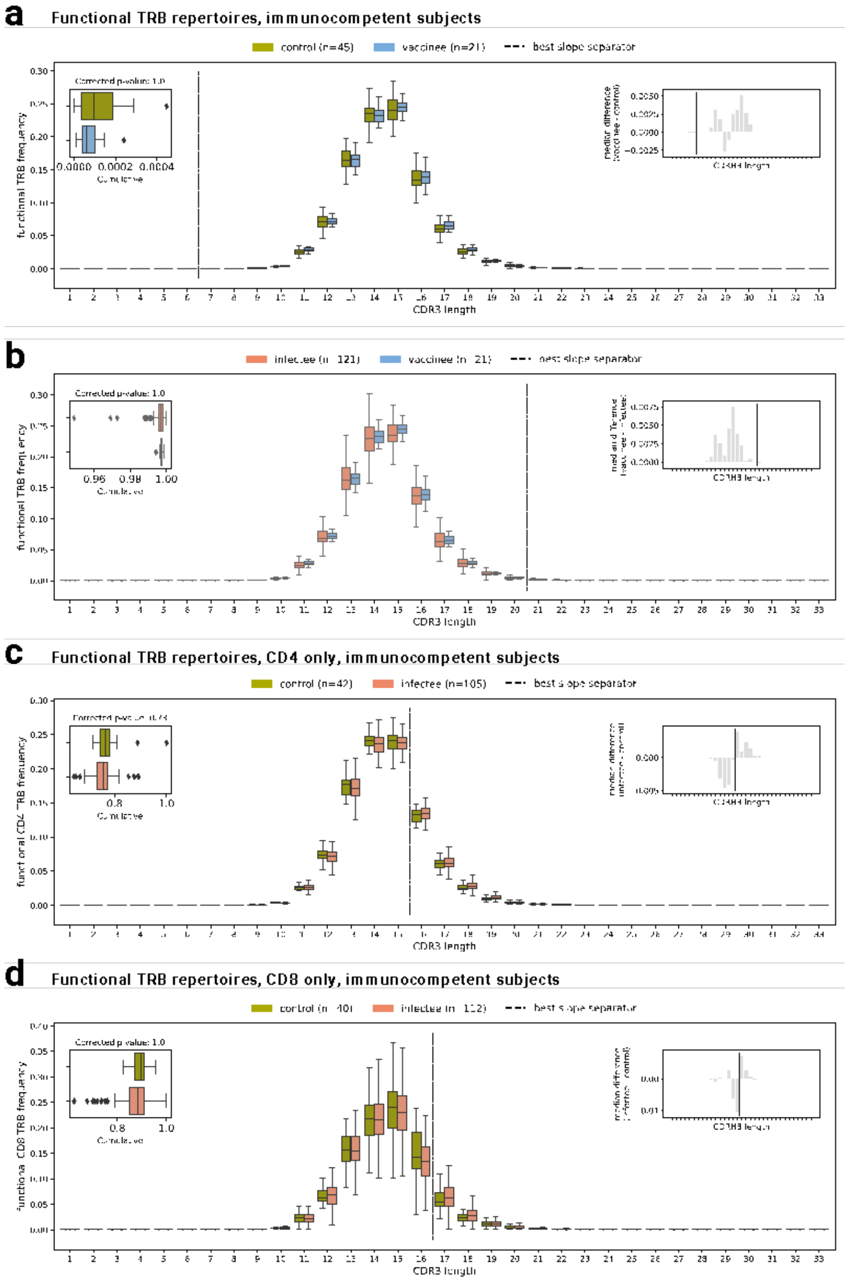

**Figure S4: CDR3 length comparison plots for functional TRD repertoires, including subsets**

Box-and-whisker plots of CDR3 length distributions (in nucleotides) in functional TCRδ genes. (a) Immunocompetent controls (olive) vs vaccinees (blue); (b) Immunocompetent infectees (salmon) vs vaccinees; (c) CD4-positive subset in immunocompetent controls vs infectees; (d) CD4-negative (predominantly CD8-positive) subset in immunocompetent controls vs infectees.

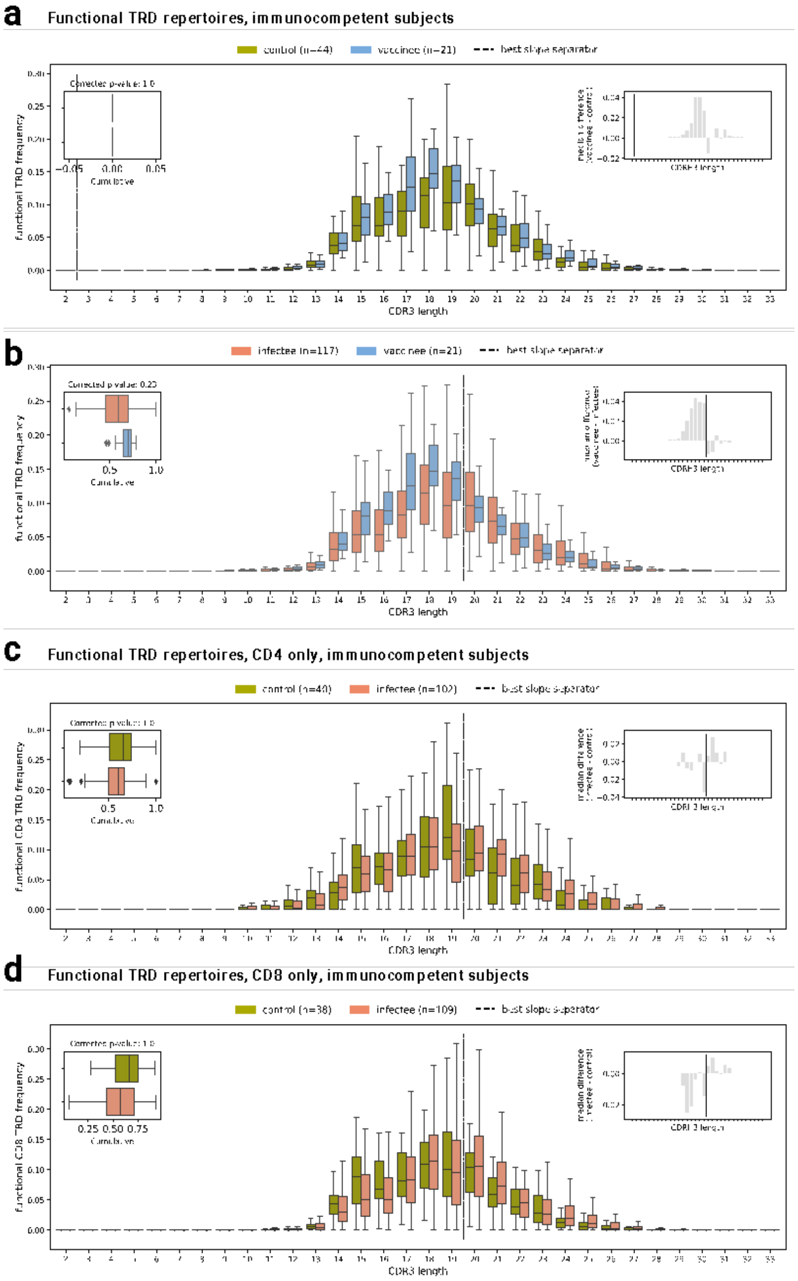

**Figure S5: CDR3 length comparison plots for non-functional IGH repertoires, including subsets**

Box-and-whisker plots of CDR3 length distributions (in nucleotides) in non-functional IGH genes. (a) Immunocompetent controls (olive) vs vaccinees (blue); (b) Immunocompetent infectees (salmon) vs vaccinees; (c) IgM-positive subset in immunocompetent controls vs infectees; (d) IgM-negative (predominantly IgG-positive) subset in immunocompetent controls vs infectees.

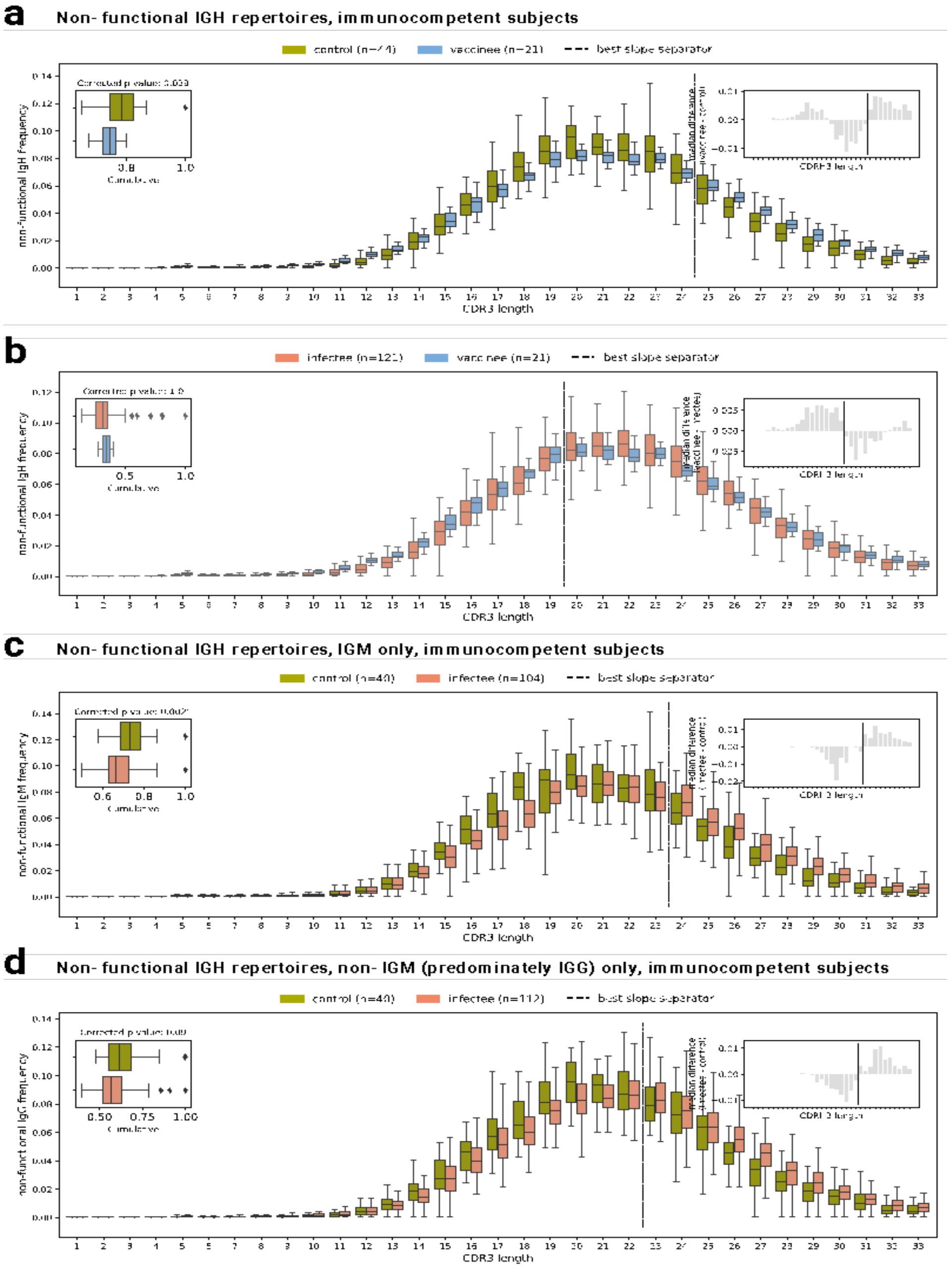

**Figure S6: CDR3 length comparison plots for non-functional TRB repertoires, including subsets**

Box-and-whisker plots of CDR3 length distributions (in nucleotides) in non-functional TCRβ genes. (a) Immunocompetent controls (olive) vs vaccinees (blue); (b) Immunocompetent infectees (salmon) vs vaccinees; (c) CD4-positive subset in immunocompetent controls vs vaccinees; (d) CD4-negative (predominantly CD8-positive) subset in immunocompetent controls vs vaccinees.

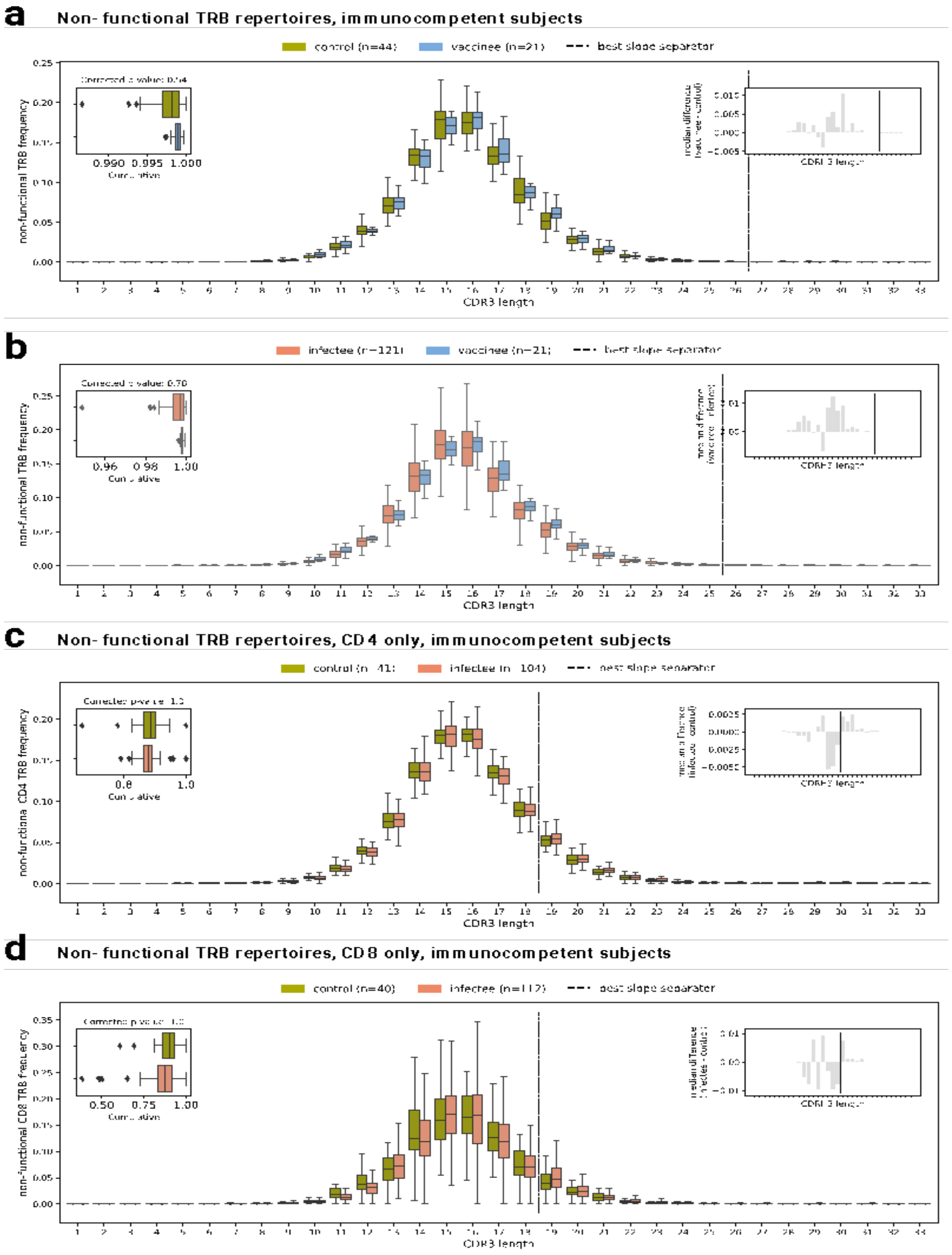

**Figure S7: CDR3 length comparison plots for non-functional TRD repertoires, including subsets**

Box-and-whisker plots of CDR3 length distributions (in nucleotides) in non-functional TCRδ genes. (a) Immunocompetent controls (olive) vs vaccinees (blue); (b) Immunocompetent infectees (salmon) vs vaccinees; (c) CD4-positive subset in immunocompetent controls vs infectees; (d) CD4-negative (i.e. predominately CD8-positive) subset in immunocompetent controls vs infectees.

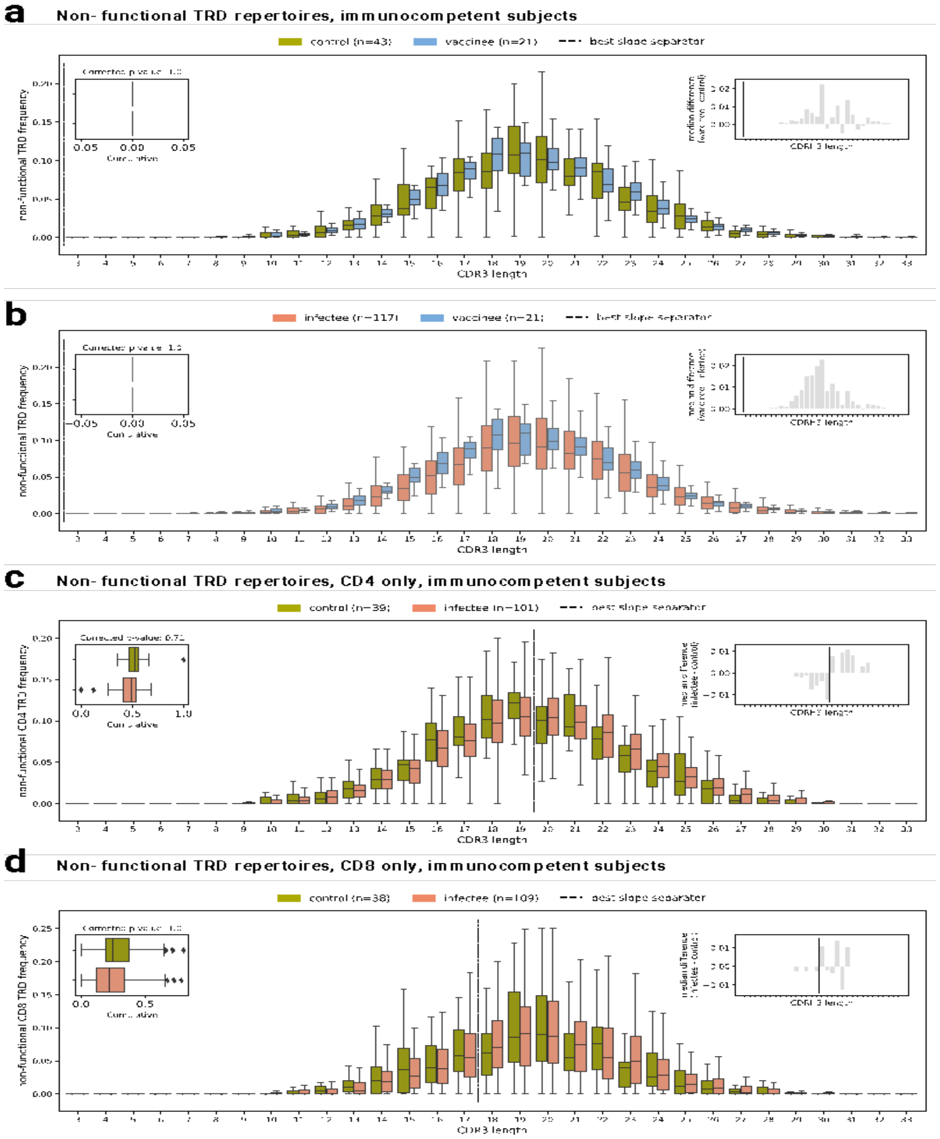

**Figure S8: Gene usage frequencies by CDR3 contribution for IGH subsets**

Box-and-whisker plots of the number of amino acids V genes and J genes contributed to the CDR3 region of IGH in the control (olive) and vaccinee (salmon) cohorts. (a) Proportion of CDR3 regions with 3 or 4 amino acids encoded by V genes in the IGM-positive subset; (b) Proportion of CDR3 regions with 5, 6, 7 or 10 amino acids encoded by J genes in the IGM-positive subset; (c) Proportion of CDR3 regions with 3 or 4 amino acids encoded by V genes in the IGM-negative subset; (d) Proportion of CDR3 regions with 5, 6, 7 or 10 amino acids encoded by J genes in the IGM-negative subset.

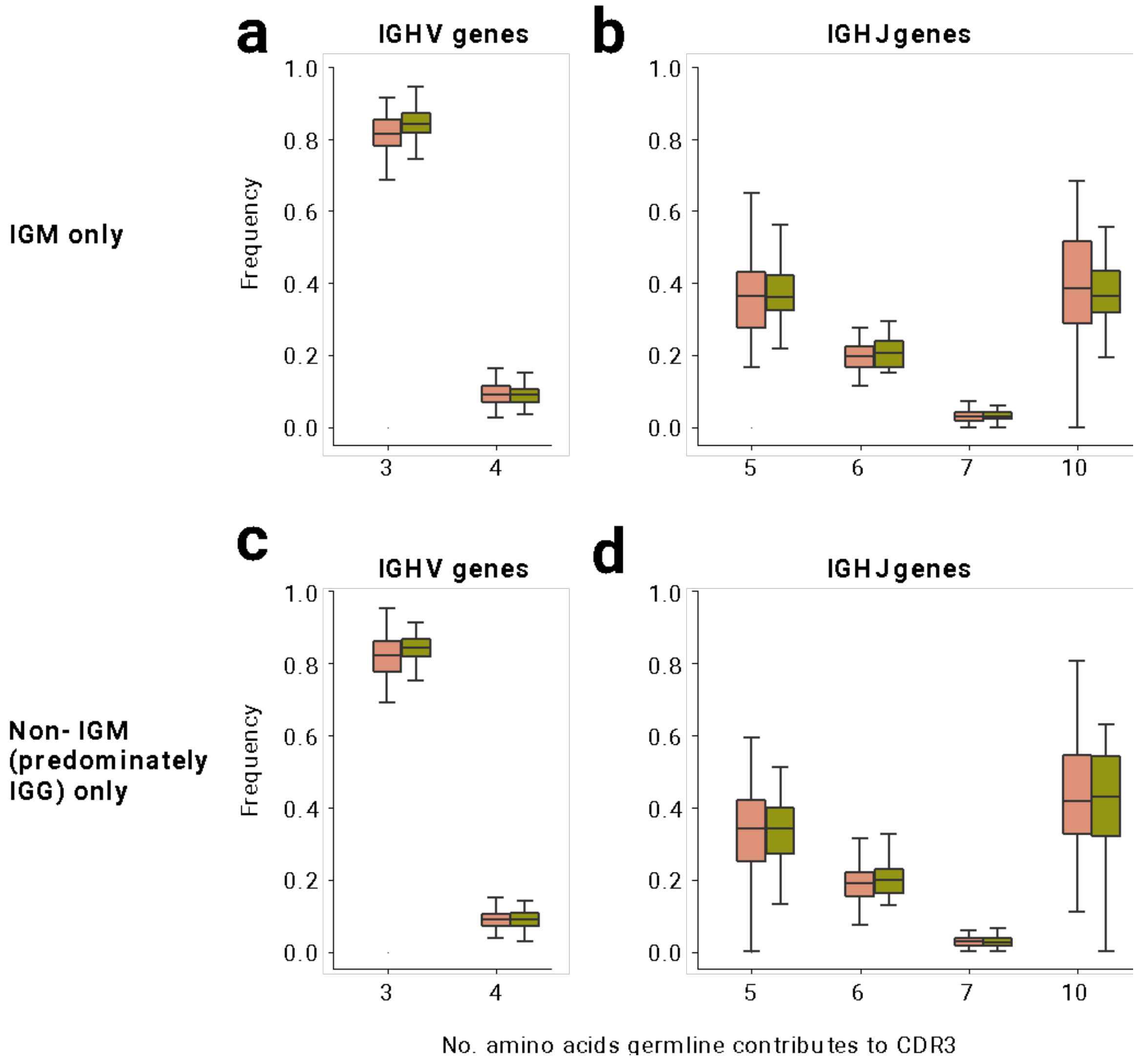

### Figure S9: TCR sequence matches vs. NAb titer

Split by cell subtype (columns) and sequence set (rows). Correlation of SARS-CoV-2 spike-specific NAb titers (BAU/mL) and fraction of database-matching sequences in the study cohort. See Methods for details.

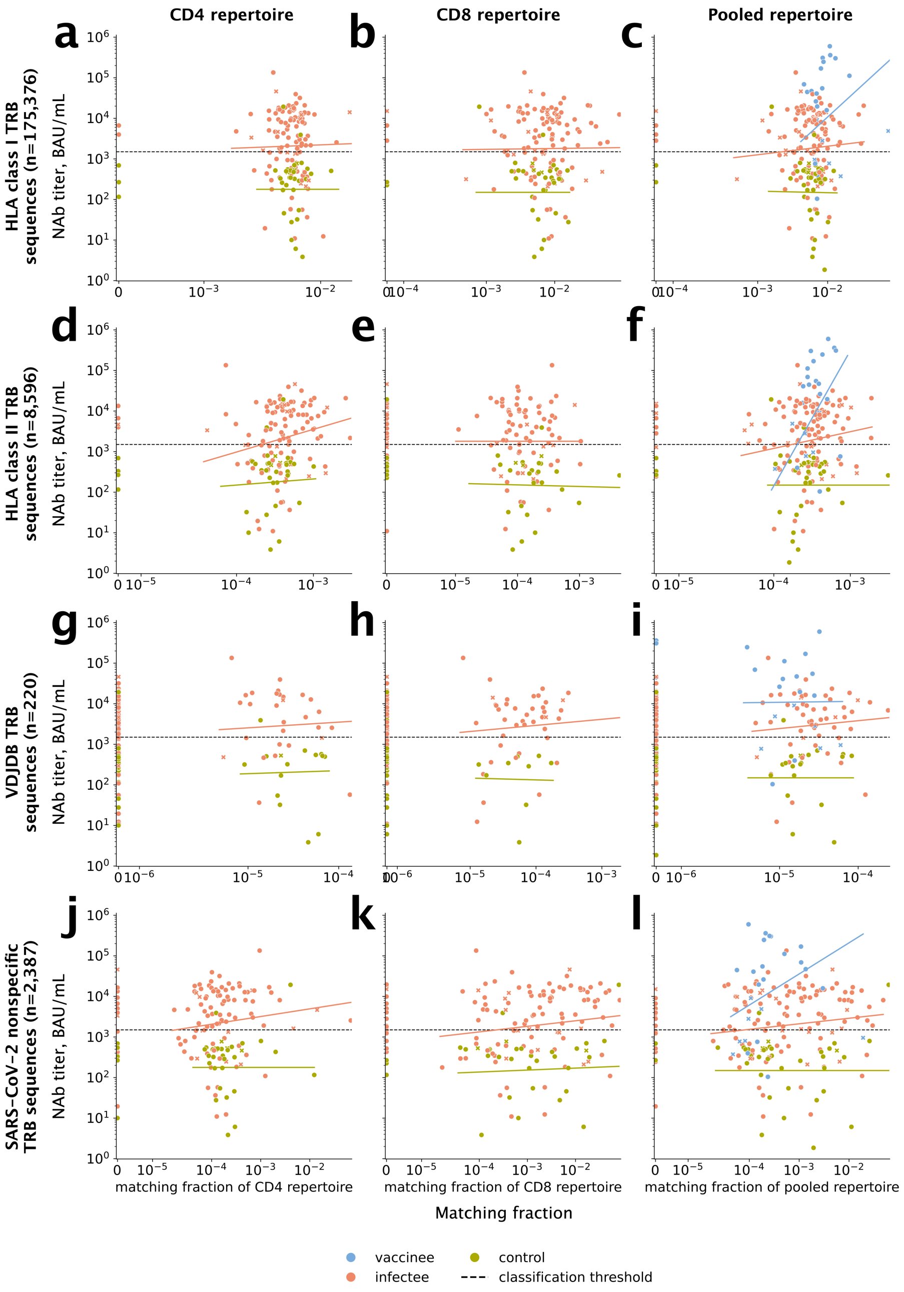

**Figure S10. Performance of fuzzy matching as a function of different tolerances**

Regardless of tolerance levels, there is an inverse correlation between repertoire sizes of the study TRB sequences and the fraction of matching reference TRB sequences. Panel (b) is reproduced from Fig. 3c for ease of comparison. See Fig. 3 legend.

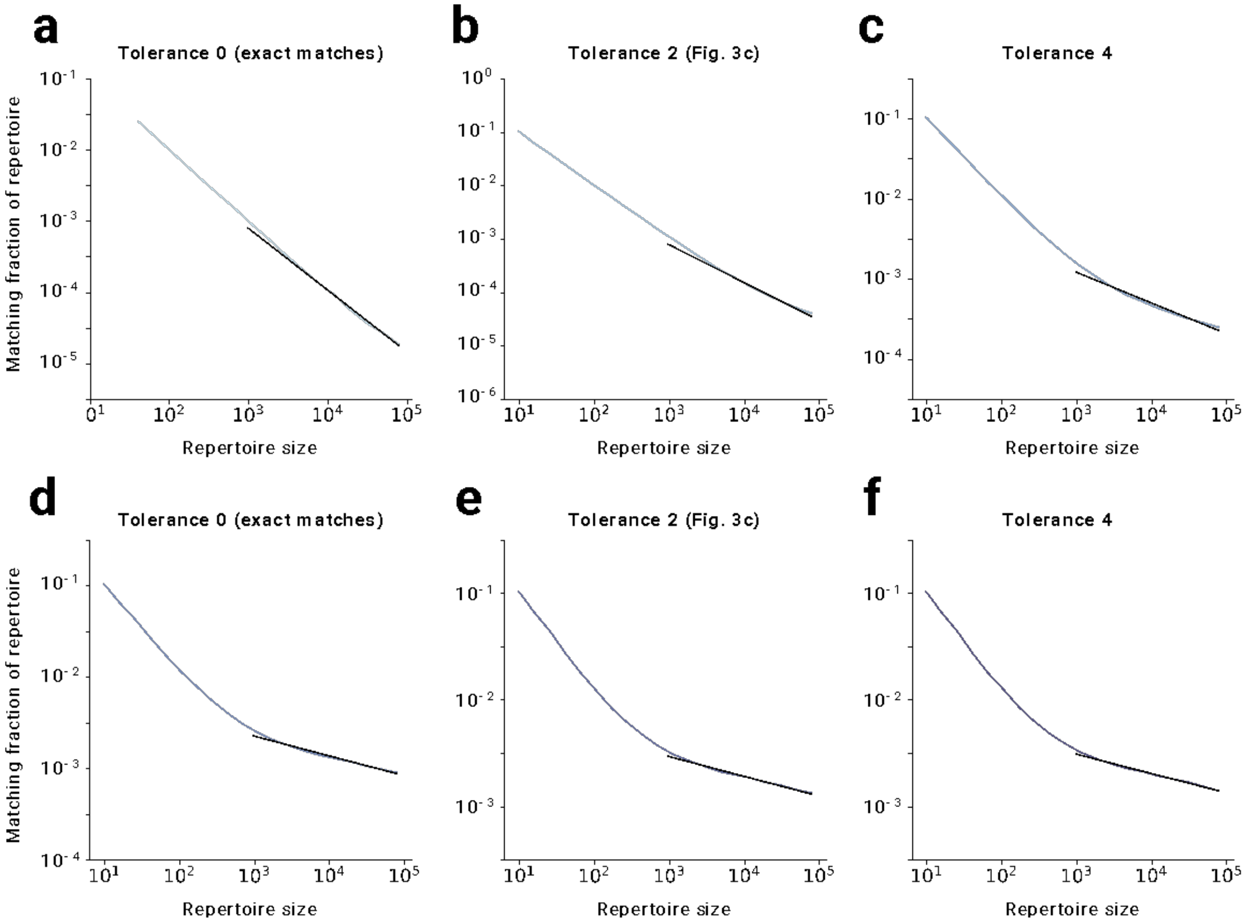

**Figure S11: Extended classifier performance**

Areas under receiver operating curves (AUROCs) of machine learning classifiers for SARS-CoV-2 exposure status per feature set and cell type, where applicable. Each boxplot corresponds to the spread of AUROCs obtained from 700 replicates (100 repeats of 7-fold cross validation). Logistic Regressions on principal components were used, in some cases preceded by feature selection. The dependent variable is SARS-CoV-2- NAb titer binned into above vs. below the manufacturer provided classification threshold for determining SARS-CoV-2 exposure.

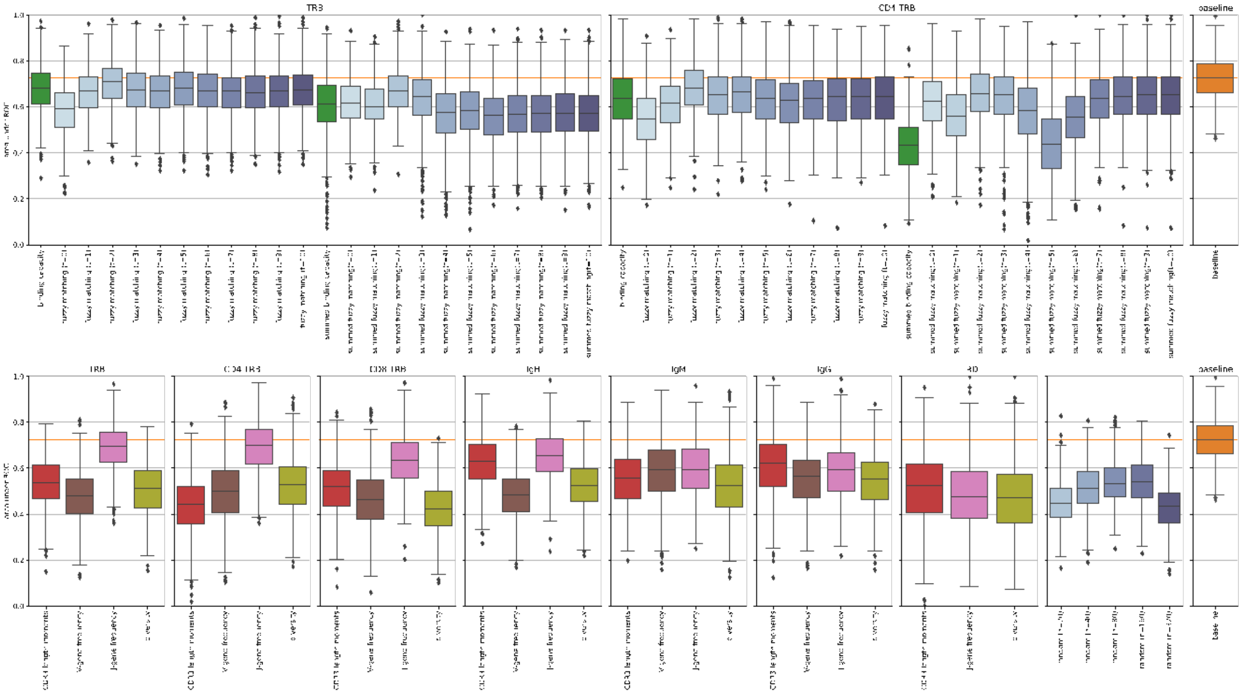

**Figure S12: Additional measures of classifier performance**

Sensitivity, precision, and specificity of machine learning classifiers for SARS-CoV-2 exposure status per feature set and cell type, where applicable. Each boxplot corresponds to the spread of values obtained from 700 replicates (100 repeats of 7-fold cross validation). Logistic Regressions on principal components were used, in some cases preceded by feature selection. The dependent variable is SARS-CoV-2-neutralizing antibody concentration binned into above or below the manufacturer provided classification threshold for determining SARS-CoV-2 exposure.

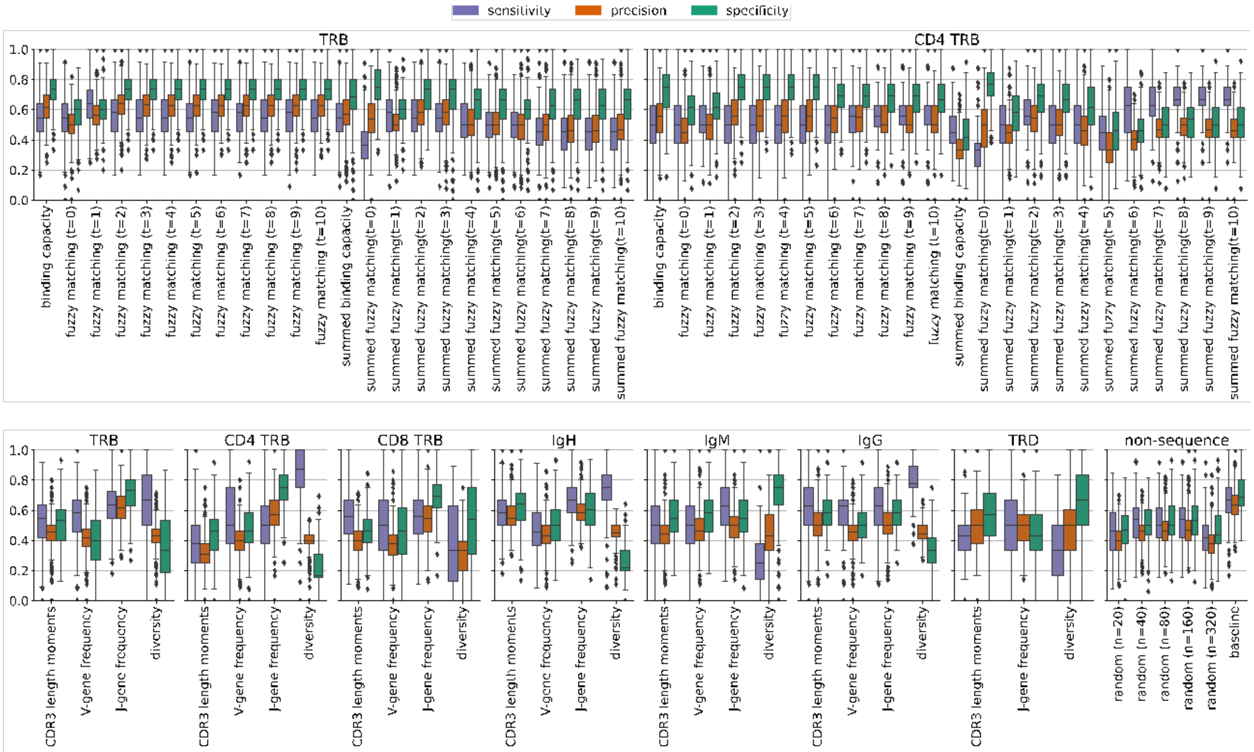

**Figure S13: Sampling times vs. NAb titers**

Times between **(a)** vaccination and sampling and **(b)** infection and sampling, with moving averages of NAb titers by ELISA, with summary box-and-whiskers plots (see also Fig. 2 legend).

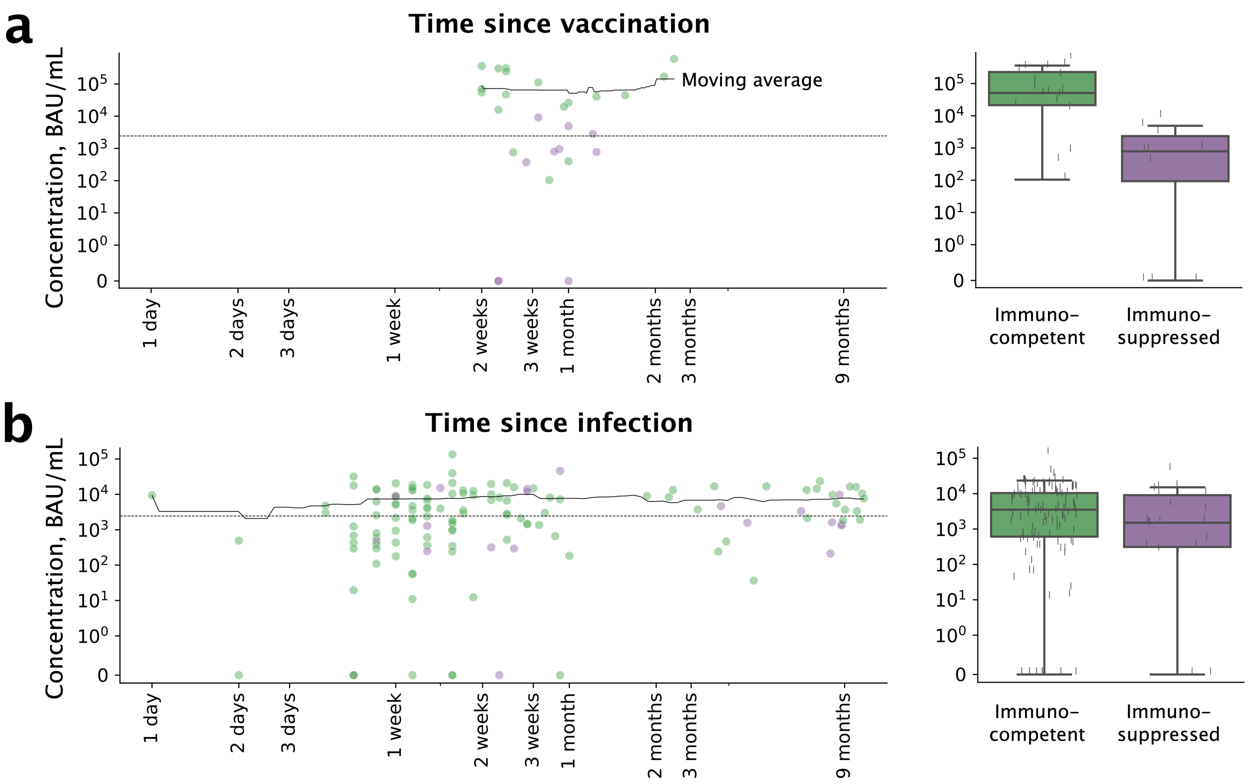

**Figure S14: Cell sorting protocol**

PBMCs were sorted in a single sorting step, as shown. All blood specimens were fractionated into plasma and PBMC. The plasma samples were used in ELISA assays to determine neutralizing SARS-CoV-2 spike-specific antibody responses. Total PBMCs were used to sequence the CDR3 region. In the vaccinee cohort, total PBMC were directly sequenced, whereas in the infectee and control cohorts, PBMC were further sorted into IGM+/CD4+ and IGM-/CD4- subfractions and the subfractions were sequenced separately.

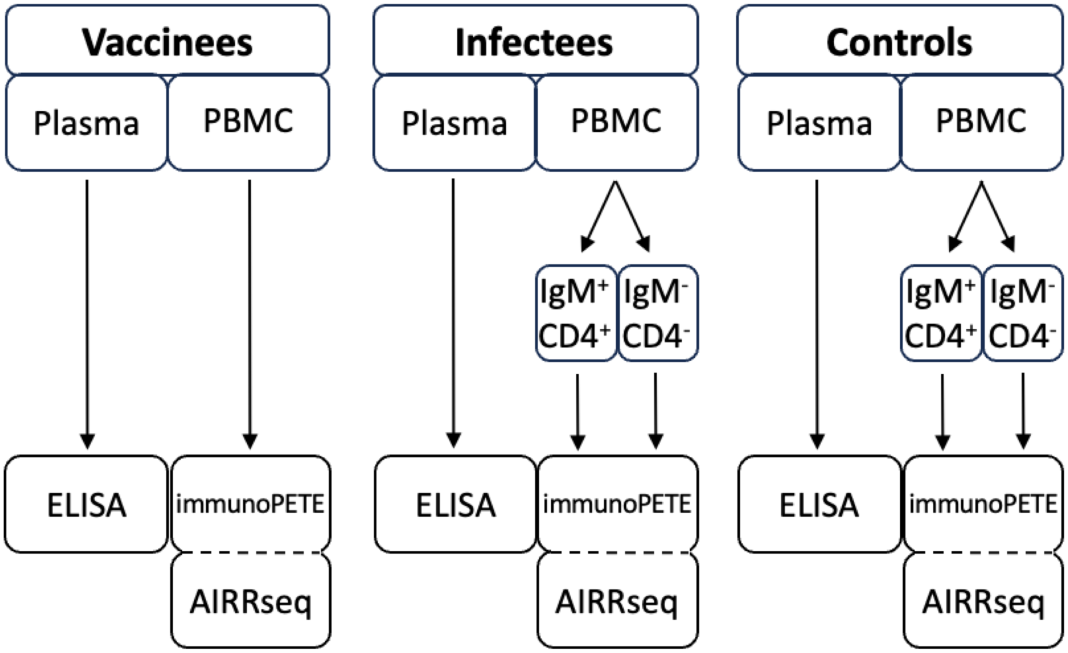

**Table S1: Demographics of study subjects**

| | | Vaccinees | Controls | Infectees | Immunosuppressed |
|---|---|---|---|---|---|
| Age (years) | $<35$ | 10 (25%) | 4 (7%) | 12 (8%) | 1 (2%) |
| | 35 - 65 | 21 (52%) | 27 (46%) | 60 (40%) | 30 (58%) |
| | $>65$ | 9 (22%) | 28 (47%) | 78 (52%) | 21 (40%) |
| Gender | M | 18 (45%) | 33 (56%) | 82 (55%) | 31 (60%) |
| | F | 22 (55%) | 26 (44%) | 68 (45%) | 21 (40%) |
| Race | unknown | 12 (30%) | 1 (2%) | 22 (15%) | 2 (4%) |
| | White | 23 (57%) | 40 (68%) | 87 (58%) | 35 (67%) |
| | Black | 4 (10%) | 11 (19%) | 29 (19%) | 11 (21%) |
| | Hispanic | 0 (0%) | 3 (5%) | 8 (5%) | 0 (0%) |
| | Asian | 1 (2%) | 4 (7%) | 4 (3%) | 3 (6%) |
| | Native American | 0 (0%) | 0 (0%) | 0 (0%) | 1 (2%) |

**Table S2: Summary of COVID-19 comorbidities**

| | | Vaccinees | Controls | Infectees | Immunosuppressed |
|---|---|---|---|---|---|
| Comorbidity Count | 0 | 9 (22%) | 0 (0%) | 4 (3%) | 0 (0%) |
| | 1 | 5 (12%) | 4 (7%) | 15 (10%) | 4 (8%) |
| | >= 2 | 26 (65%) | 55 (93%) | 131 (87%) | 48 (92%) |
| Comorbidities | Cerebrovascular disease | 4 (10%) | 8 (14%) | 20 (13%) | 8 (15%) |
| | Transplanted organ and tissue status | 17 (42%) | 3 (5%) | 15 (10%) | 31 (60%) |
| | Cancer | 11 (28%) | 20 (34%) | 25 (17%) | 22 (42%) |
| | Chronic Kidney Disease | 15 (38%) | 17 (29%) | 43 (29%) | 28 (54%) |
| | Chronic Liver Disease | 13 (32%) | 15 (25%) | 27 (18%) | 23 (44%) |
| | Chronic Lung Disease | 4 (10%) | 14 (24%) | 40 (27%) | 12 (23%) |
| | Dementia or other neurological condition | 5 (12%) | 12 (20%) | 59 (39%) | 16 (31%) |
| | Diabetes | 7 (18%) | 21 (36%) | 53 (35%) | 19 (37%) |
| | Disabilities | 2 (5%) | 7 (12%) | 20 (13%) | 5 (10%) |
| | Heart conditions | 14 (35%) | 43 (73%) | 91 (61%) | 37 (71%) |
| | HIV | 2 (5%) | 3 (5%) | 1 (1%) | 2 (4%) |
| | Mental health conditions | 7 (18%) | 26 (44%) | 59 (39%) | 21 (40%) |
| | Overweight or obesity | 22 (55%) | 41 (69%) | 105 (70%) | 40 (77%) |
| | Pregnancy or recent pregnancy | 7 (18%) | 2 (3%) | 7 (5%) | 1 (2%) |
| | Sickle cell or Thalassemia | 1 (2%) | 0 (0%) | 1 (1%) | 0 (0%) |
| | Smoking, current or former | 3 (8%) | 4 (7%) | 5 (3%) | 6 (12%) |
| | Substance abuse | 5 (12%) | 6 (10%) | 18 (12%) | 10 (19%) |
| | Tuberculosis | 0 (0%) | 0 (0%) | 2 (1%) | 1 (2%) |

**Table S3: Cohort comparisons of CDR3 length frequencies**

The statistical test used for the comparisons is described in the Methods. Multiple correction was performed using the Holm-Bonferroni method first across all p-values listed here and noted in the “initial corrected p-value” column and after follow-up tests across all p-values in this table together with the p-values in table S4 and noted in the “final corrected p-value” column. Significant corrected p-values are marked with as asterisk.

| clone type | cell type | cohort1 | cohort2 | p-value | initial corrected p-value | final corrected p-value |
|---|---|---|---|---|---|---|
| functional | pooled IgH | control | vaccinee | 0.00058 | *0.013 | *0.024 |
| functional | pooled IgH | infectee | vaccinee | 0.0001 | *0.0023 | *0.0046 |
| functional | IgM | control | infectee | 0.056 | 0.55 | 1.0 |
| functional | IgG | control | infectee | 0.027 | 0.38 | 0.84 |
| functional | pooled TCR-$\beta$ | control | vaccinee | 0.4 | 1.0 | 1.0 |
| functional | pooled TCR-$\beta$ | infectee | vaccinee | 0.52 | 1.0 | 1.0 |
| functional | CD4 TCR-$\beta$ | control | infectee | 0.022 | 0.35 | 0.73 |
| functional | CD8 TCR-$\beta$ | control | infectee | 0.086 | 0.61 | 1.0 |
| functional | pooled TCR-$\delta$ | control | vaccinee | 1.0 | 1.0 | 1.0 |
| functional | pooled TCR-$\delta$ | infectee | vaccinee | 0.0061 | 0.12 | 0.23 |
| functional | CD4 TCR-$\delta$ | control | infectee | 0.038 | 0.49 | 1.0 |
| functional | CD8 TCR-$\delta$ | control | infectee | 0.046 | 0.5 | 1.0 |
| non-functional | pooled IgH | control | vaccinee | 0.00095 | *0.02 | *0.039 |
| non-functional | pooled IgH | infectee | vaccinee | 0.042 | 0.5 | 1.0 |
| non-functional | IgM | control | infectee | 4.2e-05 | *0.001 | *0.0021 |
| non-functional | IgG | control | infectee | 0.0022 | *0.045 | 0.09 |
| non-functional | pooled TCR-$\beta$ | control | vaccinee | 0.016 | 0.28 | 0.54 |
| non-functional | pooled TCR-$\beta$ | infectee | vaccinee | 0.024 | 0.36 | 0.78 |
| non-functional | CD4 TCR-$\beta$ | control | infectee | 0.077 | 0.61 | 1.0 |
| non-functional | CD8 TCR-$\beta$ | control | infectee | 0.055 | 0.55 | 1.0 |
| non-functional | pooled TCR-$\delta$ | control | vaccinee | 1.0 | 1.0 | 1.0 |
| non-functional | pooled TCR-$\delta$ | infectee | vaccinee | 1.0 | 1.0 | 1.0 |
| non-functional | CD4 TCR-$\delta$ | control | infectee | 0.021 | 0.35 | 0.71 |
| non-functional | CD8 TCR-$\delta$ | control | infectee | 0.077 | 0.61 | 1.0 |

**Table S4: Cohort comparisons of frequencies of IGH sequences using V-/J-genes grouped by number of residues the genes contribute to the CDR3**

This table is follow-up tests to the significant CDR3 length comparisons shown in Table S3. Mann-Whitney-U tests were used to compare frequencies between cohorts. Multiple correction was performed across all p-values in this table together with the p-values in table S3 using the Holm-Bonferroni method. Significant (at α=0.05) corrected p-values are marked with an asterisk.

| clone type | gene | cell type | cohort1 | cohort2 | CDR3 residues | p-value | corrected p-value |
|---|---|---|---|---|---|---|---|
| functional | J | pooled IgH | vaccinee | infectee | 5 | 3.6e-08 | *1.9e-06 |
| functional | J | pooled IgH | vaccinee | control | 5 | 1.8e-06 | *9.4e-05 |
| functional | J | pooled IgH | vaccinee | infectee | 6 | 4.3e-05 | *0.0021 |
| functional | J | pooled IgH | vaccinee | control | 6 | 4e-06 | *0.0002 |
| functional | J | pooled IgH | vaccinee | infectee | 7 | 0.4 | 1.0 |
| functional | J | pooled IgH | vaccinee | control | 7 | 0.11 | 1.0 |
| functional | J | pooled IgH | vaccinee | infectee | 10 | 0.95 | 1.0 |
| functional | J | pooled IgH | vaccinee | control | 10 | 0.39 | 1.0 |
| functional | V | pooled IgH | vaccinee | infectee | 3 | 5.8e-05 | *0.0028$^{\dagger}$ |
| functional | V | pooled IgH | vaccinee | control | 3 | 0.00047 | *0.021$^{\dagger}$ |
| functional | V | pooled IgH | vaccinee | infectee | 4 | 5.8e-05 | *0.0028$^{\dagger}$ |
| functional | V | pooled IgH | vaccinee | control | 4 | 0.00047 | *0.021$^{\dagger}$ |
| non-functional | J | IgG | infectee | control | 5 | 0.88 | 1.0 |
| non-functional | J | IgG | infectee | control | 6 | 0.54 | 1.0 |
| non-functional | J | IgG | infectee | control | 7 | 0.65 | 1.0 |
| non-functional | J | IgG | infectee | control | 10 | 0.69 | 1.0 |
| non-functional | J | IgM | infectee | control | 5 | 0.6 | 1.0 |
| non-functional | J | IgM | infectee | control | 6 | 0.16 | 1.0 |
| non-functional | J | IgM | infectee | control | 7 | 0.79 | 1.0 |
| non-functional | J | IgM | infectee | control | 10 | 0.18 | 1.0 |
| non-functional | J | pooled IgH | vaccinee | control | 5 | 2.2e-06 | *0.00011 |
| non-functional | J | pooled IgH | vaccinee | control | 6 | 0.4 | 1.0 |
| non-functional | J | pooled IgH | vaccinee | control | 7 | 0.041 | 1.0 |
| non-functional | J | pooled IgH | vaccinee | control | 10 | 0.00052 | *0.022 |
| non-functional | V | IgG | infectee | control | 3 | 0.094$^{\dagger}$ | 1.0 |
| non-functional | V | IgG | infectee | control | 4 | 0.64$^{\dagger}$ | 1.0 |
| non-functional | V | pooled IgH | vaccinee | control | 3 | 0.0025$^{\dagger}$ | 0.098 |
| non-functional | V | pooled IgH | vaccinee | control | 4 | 0.0069$^{\dagger}$ | 0.26 |
| non-functional | V | IgM | infectee | control | 3 | 0.012$^{\dagger}$ | 0.44 |
| non-functional | V | IgM | infectee | control | 4 | 0.65$^{\dagger}$ | 1.0 |

†For productive joins, all IGH V genes contribute either 3 or 4 residues to the CDR3, so comparing frequencies of these two groups return the same p-value. Non-productive repertoires contain sequences that use V genes that are annotated as pseudogenes, which are not included in the grouping by CDR3 residue contribution but contribute to the normalizing factor when calculating usage frequency, so that p-values for the two comparisons may differ.

**Table S5: Fractions of publicly available SARS-CoV-2- and non-SARS-CoV-2-specific sequence sets matching sequences in the subject repertoires studied in this work**

| target | chain | source | matches / total (%) |
|---|---|---|---|
| SARS-CoV-2 | TRB | MIRA MHC class I | 21,719 / 175,376 (12.4%) |
| SARS-CoV-2 | TRB | MIRA MHC class II | 1,087 / 8,586 (12.7%) |
| SARS-CoV-2 | IGH | PDB/CovAbDab | 1 / 1,630 (0.06%) |
| SARS-CoV-2 | TRB | VDJDB | 45 / 220 (20.5%) |
| non-SARS-CoV-2 | IGH | PDB/CovAbDab | 0 / 340 (0.0%) |
| non-SARS-CoV-2 | TRB | VDJDB | 491 / 2,387 (20.6%) |

**Table S6: Theil-Sen estimators of linear slopes and 95% CI relating fractions of TCR repertoires matching specific clonotype sets to NAbs**

Independently fit for different clonotype sets, cell subtypes and cohorts. Regression slopes whose confidence interval does not contain 0 are marked with an asterisk.

| MHC class | subtype | cohort | slope | 95% CI |
|---|---|---|---|---|
| Type I | CD4 | control | 0.0 | (-1.8, 0.68) |
| | | infectee | 0.098 | (-0.8, 1.1) |
| | CD8 | control | 0.0 | (-0.65, 0.36) |
| | | infectee | 0.022 | (-0.32, 0.35) |
| | pooled | control | -0.038 | (-1.7, 0.19) |
| | | infectee | 0.21 | (-0.2, 0.67) |
| | | vaccinee | 1.6 | (-2.1, 5.8) |
| Type II | CD4 | control | 0.15 | (-0.13, 1.3) |
| | | infectee | *0.56 | (0.15, 1.2) |
| | CD8 | control | -0.039 | (-0.47, 0.054) |
| | | infectee | 0.0 | (-0.19, 0.2) |
| | pooled | control | 0.0 | (-0.2, 1.4) |
| | | infectee | *0.41 | (0.075, 0.81) |
| | | vaccinee | *3.3 | (0.28, 6.8) |
| VDJDB | CD4 | control | 0.075 | (-0.25, 0.48) |
| | | infectee | 0.14 | (-0.16, 0.46) |
| | CD8 | control | -0.045 | (-0.85, 0.21) |
| | | infectee | 0.15 | (-0.079, 0.41) |
| | pooled | control | 0.0 | (-0.27, 0.28) |
| | | infectee | 0.19 | (-0.023, 0.44) |
| | | vaccinee | 0.024 | (-1.1, 1.4) |
| non-COVID | CD4 | control | 0.0 | (-0.32, 0.25) |
| | | infectee | 0.19 | (-0.027, 0.45) |
| | CD8 | control | 0.048 | (-0.027, 0.36) |
| | | infectee | *0.14 | (0.0085, 0.27) |
| | pooled | control | 0.0 | (-0.1, 0.27) |
| | | infectee | *0.14 | (0.0031, 0.28) |
| | | vaccinee | 0.77 | (-0.46, 2.1) |

**Table S7: Robustness of binding capacity vs. fuzzy matching**

Slopes and 95% confidence intervals of linear mixed models fit on repertoire size vs feature values for 30 subsampled subjects' pooled TCR repertoires measured on the SARS-CoV-2 specific CD4 clonotypes from Nolan et al. (2020) using subsample sizes ≥ 1000.

| feature | slope | 95% CI |
|---|---|---|
| binding capacity | -0.11 | (-0.14, -0.093) |
| tolerance 0 | -0.87 | (-0.9, -0.84) |
| tolerance 2 | -0.71 | (-0.71, -0.7) |
| tolerance 4 | -0.38 | (-0.4, -0.36) |
| tolerance 6 | -0.22 | (-0.24, -0.2) |
| tolerance 8 | -0.18 | (-0.2, -0.16) |
| tolerance 10 | -0.18 | (-0.2, -0.16) |

### Table S8: Classifier performance

Medians and interquartile ranges for SARS-CoV-2 exposure status classifiers per cell type across the 700 replicates are given for four different performance metrics: area under receiver operating curve (AUROC), sensitivity, specificity, and precision.

| cell type | features | AUROC | sensitivity | specificity | precision |
|---|---|---|---|---|---|
| N/A | baseline | 0.72 (0.66-0.79) | 0.67 (0.58-0.75) | 0.69 (0.62-0.8 ) | 0.64 (0.57-0.7 ) |
| IgM | J-gene frequency | 0.59 (0.51-0.68) | 0.62 (0.5 -0.75) | 0.55 (0.45-0.67) | 0.5 (0.42-0.56) |
| IgM | V-gene frequency | 0.59 (0.5 -0.68) | 0.5 (0.38-0.62) | 0.58 (0.45-0.67) | 0.46 (0.4 -0.55) |
| IgM | CDR3 length moments | 0.56 (0.47-0.64) | 0.5 (0.38-0.62) | 0.55 (0.45-0.67) | 0.44 (0.38-0.5 ) |
| IgM | diversity | 0.52 (0.43-0.61) | 0.25 (0.14-0.38) | 0.75 (0.64-0.83) | 0.43 (0.33-0.57) |
| IgG | J-gene frequency | 0.59 (0.5 -0.67) | 0.62 (0.5 -0.75) | 0.58 (0.5 -0.67) | 0.5 (0.44-0.58) |
| IgG | V-gene frequency | 0.56 (0.47-0.64) | 0.62 (0.5 -0.67) | 0.5 (0.42-0.58) | 0.45 (0.4 -0.5 ) |
| IgG | CDR3 length moments | 0.62 (0.52-0.7 ) | 0.62 (0.5 -0.75) | 0.58 (0.5 -0.67) | 0.5 (0.43-0.58) |
| IgG | diversity | 0.55 (0.46-0.62) | 0.78 (0.75-0.89) | 0.33 (0.25-0.42) | 0.44 (0.41-0.5 ) |
| pooled IgH | J-gene frequency | 0.65 (0.58-0.73) | 0.67 (0.58-0.75) | 0.6 (0.5 -0.71) | 0.58 (0.53-0.64) |
| pooled IgH | V-gene frequency | 0.48 (0.41-0.55) | 0.45 (0.36-0.58) | 0.5 (0.4 -0.6 ) | 0.43 (0.38-0.5 ) |
| pooled IgH | CDR3 length moments | 0.63 (0.55-0.7 ) | 0.58 (0.5 -0.67) | 0.64 (0.53-0.71) | 0.55 (0.5 -0.62) |
| pooled IgH | diversity | 0.53 (0.45-0.6 ) | 0.75 (0.67-0.83) | 0.21 (0.2 -0.33) | 0.45 (0.42-0.5 ) |
| CD8 TCR-$\beta$ | J-gene frequency | 0.63 (0.56-0.71) | 0.56 (0.44-0.67) | 0.69 (0.62-0.77) | 0.55 (0.45-0.62) |
| CD8 TCR-$\beta$ | V-gene frequency | 0.46 (0.38-0.55) | 0.5 (0.33-0.67) | 0.46 (0.31-0.62) | 0.38 (0.3 -0.44) |
| CD8 TCR-$\beta$ | CDR3 length moments | 0.52 (0.43-0.59) | 0.56 (0.44-0.67) | 0.46 (0.38-0.54) | 0.4 (0.33-0.46) |
| CD8 TCR-$\beta$ | diversity | 0.42 (0.35-0.5 ) | 0.33 (0.12-0.62) | 0.54 (0.31-0.75) | 0.33 (0.2 -0.39) |
| pooled TCR-$\delta$ | J-gene frequency | 0.48 (0.38-0.58) | 0.5 (0.33-0.67) | 0.43 (0.33-0.57) | 0.5 (0.4 -0.57) |
| pooled TCR-$\delta$ | CDR3 length moments | 0.52 (0.4 -0.62) | 0.43 (0.33-0.5 ) | 0.57 (0.43-0.71) | 0.5 (0.38-0.6 ) |
| pooled TCR-$\delta$ | diversity | 0.47 (0.36-0.57) | 0.33 (0.17-0.5 ) | 0.67 (0.5 -0.83) | 0.5 (0.33-0.6 ) |
| CD4 TCR-$\beta$ | J-gene frequency | 0.7 (0.62-0.77) | 0.5 (0.38-0.62) | 0.75 (0.67-0.83) | 0.57 (0.5 -0.67) |
| CD4 TCR-$\beta$ | V-gene frequency | 0.5 (0.4 -0.59) | 0.5 (0.38-0.75) | 0.46 (0.38-0.58) | 0.4 (0.33-0.46) |
| CD4 TCR-$\beta$ | CDR3 length moments | 0.44 (0.36-0.52) | 0.38 (0.25-0.5 ) | 0.46 (0.33-0.54) | 0.31 (0.25-0.38) |
| CD4 TCR-$\beta$ | diversity | 0.53 (0.44-0.61) | 0.88 (0.75-1.0 ) | 0.17 (0.15-0.31) | 0.4 (0.38-0.44) |
| CD4 TCR-$\beta$ | binding capacity | 0.64 (0.55-0.72) | 0.5 (0.38-0.62) | 0.75 (0.62-0.83) | 0.56 (0.44-0.64) |
| CD4 TCR-$\beta$ | fuzzy matching (t=0) | 0.55 (0.46-0.63) | 0.5 (0.38-0.62) | 0.62 (0.5 -0.69) | 0.44 (0.38-0.55) |
| CD4 TCR-$\beta$ | fuzzy matching (t=1) | 0.62 (0.53-0.69) | 0.5 (0.44-0.62) | 0.62 (0.54-0.71) | 0.5 (0.4 -0.57) |
| CD4 TCR-$\beta$ | fuzzy matching (t=2) | 0.68 (0.61-0.76) | 0.5 (0.38-0.62) | 0.75 (0.67-0.83) | 0.56 (0.5 -0.67) |
| CD4 TCR-$\beta$ | fuzzy matching (t=3) | 0.65 (0.57-0.73) | 0.5 (0.38-0.62) | 0.75 (0.67-0.83) | 0.56 (0.45-0.67) |
| CD4 TCR-$\beta$ | fuzzy matching (t=4) | 0.66 (0.57-0.73) | 0.5 (0.38-0.62) | 0.75 (0.67-0.83) | 0.56 (0.44-0.67) |
| CD4 TCR-$\beta$ | fuzzy matching (t=5) | 0.63 (0.55-0.72) | 0.5 (0.38-0.62) | 0.75 (0.67-0.83) | 0.56 (0.45-0.67) |
| CD4 TCR-$\beta$ | fuzzy matching (t=6) | 0.63 (0.53-0.7 ) | 0.5 (0.38-0.62) | 0.69 (0.62-0.77) | 0.54 (0.44-0.62) |
| CD4 TCR-$\beta$ | fuzzy matching (t=7) | 0.64 (0.55-0.71) | 0.56 (0.44-0.62) | 0.69 (0.62-0.77) | 0.55 (0.45-0.62) |
| CD4 TCR-$\beta$ | fuzzy matching (t=8) | 0.64 (0.54-0.72) | 0.56 (0.49-0.62) | 0.69 (0.58-0.77) | 0.5 (0.44-0.62) |
| CD4 TCR-$\beta$ | fuzzy matching (t=9) | 0.64 (0.55-0.72) | 0.56 (0.5 -0.62) | 0.69 (0.58-0.77) | 0.5 (0.44-0.62) |
| CD4 TCR-$\beta$ | fuzzy matching (t=10) | 0.64 (0.55-0.73) | 0.62 (0.5 -0.62) | 0.67 (0.58-0.77) | 0.5 (0.44-0.62) |
| CD4 TCR-$\beta$ | summed binding capacity | 0.43 (0.35-0.51) | 0.44 (0.38-0.56) | 0.42 (0.33-0.54) | 0.33 (0.27-0.4 ) |
| CD4 TCR-$\beta$ | summed fuzzy matching(t=0) | 0.62 (0.54-0.71) | 0.33 (0.22-0.38) | 0.77 (0.69-0.85) | 0.5 (0.35-0.6 ) |
| CD4 TCR-$\beta$ | summed fuzzy matching(t=1) | 0.56 (0.47-0.65) | 0.5 (0.38-0.62) | 0.58 (0.46-0.69) | 0.44 (0.38-0.5 ) |
| CD4 TCR-$\beta$ | summed fuzzy matching(t=2) | 0.66 (0.58-0.74) | 0.56 (0.5 -0.67) | 0.69 (0.58-0.77) | 0.55 (0.45-0.62) |
| CD4 TCR-$\beta$ | summed fuzzy matching(t=3) | 0.65 (0.57-0.73) | 0.5 (0.38-0.62) | 0.67 (0.58-0.77) | 0.5 (0.43-0.6 ) |
| CD4 TCR-$\beta$ | summed fuzzy matching(t=4) | 0.58 (0.48-0.68) | 0.5 (0.38-0.62) | 0.62 (0.5 -0.75) | 0.46 (0.36-0.56) |
| CD4 TCR-$\beta$ | summed fuzzy matching(t=5) | 0.44 (0.33-0.55) | 0.44 (0.33-0.56) | 0.46 (0.33-0.58) | 0.33 (0.25-0.44) |
| CD4 TCR-$\beta$ | summed fuzzy matching(t=6) | 0.55 (0.46-0.64) | 0.62 (0.44-0.75) | 0.46 (0.38-0.54) | 0.4 (0.33-0.46) |
| CD4 TCR-$\beta$ | summed fuzzy matching(t=7) | 0.63 (0.55-0.72) | 0.62 (0.56-0.75) | 0.5 (0.42-0.62) | 0.47 (0.42-0.54) |
| CD4 TCR-$\beta$ | summed fuzzy matching(t=8) | 0.65 (0.57-0.73) | 0.67 (0.62-0.75) | 0.54 (0.42-0.62) | 0.5 (0.43-0.55) |
| CD4 TCR-$\beta$ | summed fuzzy matching(t=9) | 0.65 (0.57-0.73) | 0.67 (0.62-0.75) | 0.5 (0.42-0.62) | 0.47 (0.42-0.54) |
| CD4 TCR-$\beta$ | summed fuzzy matching(t=10) | 0.65 (0.57-0.73) | 0.67 (0.62-0.75) | 0.5 (0.42-0.62) | 0.46 (0.42-0.54) |
| pooled TCR-$\beta$ | J-gene frequency | 0.69 (0.62-0.76) | 0.64 (0.55-0.73) | 0.73 (0.62-0.8 ) | 0.62 (0.55-0.69) |
| pooled TCR-$\beta$ | V-gene frequency | 0.48 (0.4 -0.55) | 0.58 (0.45-0.67) | 0.4 (0.27-0.53) | 0.42 (0.36-0.47) |
| pooled TCR-$\beta$ | CDR3 length moments | 0.53 (0.46-0.61) | 0.55 (0.42-0.64) | 0.53 (0.4 -0.6 ) | 0.45 (0.4 -0.5 ) |
| pooled TCR-$\beta$ | diversity | 0.51 (0.42-0.59) | 0.67 (0.5 -0.83) | 0.33 (0.19-0.53) | 0.43 (0.38-0.47) |
| pooled TCR-$\beta$ | binding capacity | 0.68 (0.61-0.74) | 0.55 (0.45-0.64) | 0.73 (0.67-0.8 ) | 0.62 (0.55-0.7 ) |
| pooled TCR-$\beta$ | fuzzy matching (t=0) | 0.59 (0.51-0.66) | 0.55 (0.45-0.64) | 0.6 (0.5 -0.67) | 0.5 (0.44-0.57) |
| pooled TCR-$\beta$ | fuzzy matching (t=1) | 0.67 (0.59-0.73) | 0.64 (0.55-0.73) | 0.6 (0.53-0.67) | 0.56 (0.5 -0.62) |
| pooled TCR-$\beta$ | fuzzy matching (t=2) | 0.71 (0.64-0.77) | 0.58 (0.45-0.67) | 0.73 (0.67-0.8 ) | 0.64 (0.57-0.7 ) |
| pooled TCR-$\beta$ | fuzzy matching (t=3) | 0.67 (0.6 -0.74) | 0.55 (0.45-0.67) | 0.73 (0.67-0.8 ) | 0.63 (0.56-0.7 ) |
| pooled TCR-$\beta$ | fuzzy matching (t=4) | 0.67 (0.59-0.73) | 0.55 (0.45-0.67) | 0.73 (0.67-0.8 ) | 0.62 (0.56-0.7 ) |
| pooled TCR-$\beta$ | fuzzy matching (t=5) | 0.68 (0.61-0.75) | 0.55 (0.45-0.64) | 0.73 (0.67-0.8 ) | 0.62 (0.56-0.7 ) |
| pooled TCR-$\beta$ | fuzzy matching (t=6) | 0.67 (0.6 -0.74) | 0.58 (0.45-0.67) | 0.73 (0.67-0.8 ) | 0.62 (0.56-0.7 ) |
| pooled TCR-$\beta$ | fuzzy matching (t=7) | 0.67 (0.59-0.73) | 0.58 (0.45-0.64) | 0.73 (0.67-0.8 ) | 0.62 (0.56-0.7 ) |
| pooled TCR-$\beta$ | fuzzy matching (t=8) | 0.66 (0.59-0.73) | 0.55 (0.45-0.67) | 0.73 (0.67-0.8 ) | 0.62 (0.56-0.7 ) |
| pooled TCR-$\beta$ | fuzzy matching (t=9) | 0.67 (0.6 -0.73) | 0.58 (0.45-0.67) | 0.73 (0.67-0.8 ) | 0.62 (0.56-0.7 ) |
| pooled TCR-$\beta$ | fuzzy matching (t=10) | 0.67 (0.61-0.74) | 0.55 (0.45-0.67) | 0.73 (0.67-0.8 ) | 0.62 (0.56-0.7 ) |
| pooled TCR-$\beta$ | summed binding capacity | 0.61 (0.53-0.69) | 0.55 (0.45-0.64) | 0.69 (0.6 -0.8 ) | 0.57 (0.5 -0.67) |
| pooled TCR-$\beta$ | summed fuzzy matching(t=0) | 0.62 (0.55-0.69) | 0.36 (0.27-0.45) | 0.75 (0.67-0.87) | 0.54 (0.44-0.64) |
| pooled TCR-$\beta$ | summed fuzzy matching(t=1) | 0.6 (0.55-0.68) | 0.58 (0.45-0.67) | 0.6 (0.53-0.67) | 0.5 (0.46-0.57) |
| pooled TCR-$\beta$ | summed fuzzy matching(t=2) | 0.67 (0.6 -0.73) | 0.55 (0.45-0.67) | 0.73 (0.6 -0.8 ) | 0.58 (0.5 -0.67) |
| pooled TCR-$\beta$ | summed fuzzy matching(t=3) | 0.64 (0.56-0.72) | 0.55 (0.45-0.64) | 0.73 (0.6 -0.8 ) | 0.58 (0.5 -0.67) |
| pooled TCR-$\beta$ | summed fuzzy matching(t=4) | 0.57 (0.48-0.65) | 0.5 (0.42-0.64) | 0.67 (0.53-0.73) | 0.5 (0.43-0.6 ) |
| pooled TCR-$\beta$ | summed fuzzy matching(t=5) | 0.58 (0.5 -0.67) | 0.5 (0.42-0.58) | 0.67 (0.53-0.73) | 0.5 (0.44-0.58) |
| pooled TCR-$\beta$ | summed fuzzy matching(t=6) | 0.56 (0.48-0.64) | 0.5 (0.42-0.58) | 0.6 (0.53-0.67) | 0.5 (0.4 -0.57) |
| pooled TCR-$\beta$ | summed fuzzy matching(t=7) | 0.57 (0.49-0.65) | 0.45 (0.36-0.55) | 0.62 (0.53-0.73) | 0.5 (0.4 -0.57) |
| pooled TCR-$\beta$ | summed fuzzy matching(t=8) | 0.57 (0.49-0.65) | 0.45 (0.33-0.55) | 0.67 (0.53-0.73) | 0.46 (0.38-0.56) |
| pooled TCR-$\beta$ | summed fuzzy matching(t=9) | 0.57 (0.49-0.65) | 0.45 (0.33-0.55) | 0.62 (0.53-0.73) | 0.46 (0.38-0.56) |
| pooled TCR-$\beta$ | summed fuzzy matching(t=10) | 0.57 (0.49-0.65) | 0.45 (0.33-0.55) | 0.67 (0.53-0.73) | 0.47 (0.4 -0.57) |
| N/A | random (n=20) | 0.45 (0.39-0.51) | 0.46 (0.33-0.58) | 0.47 (0.38-0.56) | 0.4 (0.33-0.46) |
| N/A | random (n=40) | 0.51 (0.45-0.58) | 0.54 (0.42-0.62) | 0.5 (0.44-0.6 ) | 0.46 (0.4 -0.5 ) |
| N/A | random (n=80) | 0.53 (0.47-0.6 ) | 0.5 (0.42-0.62) | 0.5 (0.44-0.62) | 0.46 (0.4 -0.53) |
| N/A | random (n=160) | 0.54 (0.47-0.61) | 0.54 (0.42-0.62) | 0.53 (0.44-0.62) | 0.47 (0.4 -0.53) |
| N/A | random (n=320) | 0.43 (0.36-0.49) | 0.42 (0.33-0.5 ) | 0.44 (0.38-0.56) | 0.38 (0.31-0.44) |
